# Temporal Dynamics of the Tomato Rhizosphere Microbiome in Response to Synthetic Communities of Plant Growth-Promoting Rhizobacteria

**DOI:** 10.1101/2024.12.21.629910

**Authors:** Daniele Nicotra, Alexandros Mosca, Giulio Dimaria, Matilde Tessitori, Ramesh Raju Vetukuri, Vittoria Catara

## Abstract

In sustainable agriculture, the application of microorganisms to soil is a widely adopted strategy aimed at enhancing soil microbiome functionality, restoring fertility, and recovering biodiversity diminished by intensive farming. While introducing individual beneficial microbes often encounters issues with establishment and persistence, multispecies microbial consortia offer a more robust alternative, providing complementary functions that enhance both their resilience and effectiveness. To test this approach, we designed three synthetic bacterial communities (SynComs), each composed of varying combinations of ten bacterial endophytes previously shown to promote plant growth and exert biocontrol effects. The SynComs were designed with ascending levels of richness (4, 6, and 10 members) and diversity (ranging from 3 to 6 bacterial genera). In growth chamber trials with tomatoes, the SynComs not only promoted plant growth but also induced significant shifts in the rhizosphere bacterial communities, primarily affecting less abundant taxa. The SynComs MIX2 and MIX3, which included *Pseudomonas* species, exhibited the greatest impact on both plant growth enhancement and shifts within the resident microbial community. Monitoring of the introduced strains over time demonstrated that most bioinoculants successfully established although at very low concentration in the rhizosphere.

## 1. Introduction

Microorganisms associated with plant roots provide a range of fundamental services for plant growth and health. Various microbiome members promote plant growth directly by producing hormones and facilitating nutrient mineralization, and indirectly through biocontrol of pathogens and induction of plant resistance (Berendsen *et al*., 2012; Compant *et al*., 2024; Mendes *et al*., 2013; Trivedi *et al*., 2021; Vandenkoornhuyse *et al*., 2015).

In sustainable agriculture, the application of microorganisms to the soil is a widely adopted practice (Afridi *et al*., 2022; Compant *et al*., 2019; Trivedi *et al*., 2021). The addition of exogenous microorganisms aims to enhance microbiome-associated multifunctionality, restoring soil fertility and biodiversity lost due to intensive farming (Delgado-Baquerizo *et al*., 2016; Upadhyay *et al*., 2023; Wubs *et al*., 2016). However, these effects can be unstable and transient if the introduced species fail to establish and survive at sufficient densities to function effectively within the natural microbiome (Compant *et al*., 2019; Haskett *et al*., 2021; Trivedi *et al*., 2020). Indeed, competition and antagonism from the resident microbiome can severely limit inoculant colonization and function in the rhizosphere (Shi *et al*., 2016; Thakur and Geisen, 2019; Trivedi *et al*., 2020). While the introduction of individual beneficial microbes often faces challenges in establishment and persistence, multispecies microbial consortia (i.e. combinations of beneficial microorganisms each playing unique roles in plant health) present a more resilient solution by offering complementary functions that enhance both survival and efficacy (de Souza *et al*., 2020; Großkopf and Soyer, 2014; Martins *et al*., 2023; Singh *et al*., 2023). The increased diversity in consortium members can also trigger the expression of previously “silent” traits, such as secondary metabolite production (Jousset *et al*., 2014; Tyc *et al*., 2014) or enzymatic activities (Weidner *et al*., 2015), enhancing functional diversity and redundancy at the consortium level. Diverse microbial communities can occupy different ecological niches and adapt to varying conditions, leading to better competition with the existing microbial communities in the rhizosphere (Hassani *et al*., 2018; Hu *et al*., 2017).

Since introduced microorganisms interact with the resident microflora, in addition to the direct effect due to the contribution of functions of individual consortium members, indirect effects can be triggered, mediated by the change in the diversity, composition and functioning of the resident microbiome (Castro-Sowinski *et al*., 2007; Vuolo *et al*., 2022). These shifts not only affect microbial competition but also modulate plant nutrient uptake efficiency, alter hormone signaling pathways, and trigger enhanced immune responses, all of which contribute to improved plant resilience (Hu *et al*., 2021; Vuolo *et al*., 2022).

Studies have shown that adding “foreign” members to the endogenous microbiome can significantly impact existing communities and affect plant growth and health (Chang *et al*., 2024; Hu *et al*., 2021; Sharma *et al*., 2022; Z. Wang *et al*., 2021; Zhang *et al*., 2022; Zheng *et al*., 2020). In some cases, these effects are transient (Buddrus-Schiemann *et al*., 2010; Qiao *et al*., 2017; Yin *et al*., 2013). Such impacts can arise from direct microbial interactions, such as competition or antagonism (Hassani *et al*., 2018), which can shift the balance of dominant and rare taxa (Hu *et al*., 2021), or from plant-mediated changes in root exudation profiles (Gu *et al*., 2016; Yuan *et al*., 2018). The extent of these alterations may correlate with the diversity and richness of the consortium, with greater diversity and richness leading to more significant changes (Rivett *et al*., 2018).

Our previous work focused on the individual effects of ten bacterial strains, each of which demonstrated promising growth-promotion and biocontrol properties (Nicotra *et al*., 2024). However, we had not previously explored the potential synergies between these strains when combined into a consortium.

The strains were isolated from tomato seeds and roots and were selected using a top-down approach based on the tomato core microbiome (Anzalone *et al*., 2022; Nicotra *et al*., 2024). Since antimicrobial activity was not used as a criterion for prescreening, strains from the less-studied genera *Leclercia*, *Chryseobacterium*, *Glutamicibacter*, and *Paenarthrobacter*, along with the well-studied biocontrol agents *Pseudomonas* and *Bacillus*, were included in the set. Both the efficacy demonstrated in plant assays and the genome traits linked to plant-microbiome interactions encouraged us to explore the fundamental interactions and potential synergies when combining these previously studied single strains.

The three small consortia (hereafter referred to as Syncom) assembled for this purpose exhibited plant growth-promoting (PGP) activity, though to varying extents. They significantly impacted the rhizosphere bacterial communities as early as one week after treatment, with most of the shifts occurring in less abundant taxa. The monitoring of the inoculants in the microbiome was attemped and eight ASVs with ≥ 99.75% sequence identity to the inoculated strains were tracked over time.

## 2. Materials and Methods

### 2.1 Bacterial strains used in this study

The ten bacteria used in this study were selected as bioinoculants amongst the core microbiome genera of tomato plants in the cultivation chain (Nicotra *et al*., 2024). The strains used belonged to six genera, namely *Bacillus*, *Pseudomonas, Glutamicibacter, Paenarthrobacter, Chryseobacterium* and *Leclercia.* Further details about the strains used and their isolation source can be found in Table 1. The genomes of the bacterial strains are available under BioProject ID: PRJNA1096641.

**Table 1.** Bacterial strains used in this study.

| Strain | Source | Genome ID |
| --- | --- | --- |
| *Pseudomonas salmasensis* POE54 | Nursery (peat) | SAMN40757903 |
| *P. simiae* POE78A | Nursery (peat) | SAMN40757904 |
| *Chryseobacterium* sp. POE47 | Nursery (peat) | SAMN40757905 |
| *Leclercia* sp. S52 | Seed | SAMN40757906 |
| *Bacillus velezensis* PSE31B | Soil | SAMN40757907 |
| *B. velezensis* PFE42 | Coconut fiber | SAMN40757908 |
| *B. velezensis* PFE11 | Coconut fiber | SAMN40757909 |
| *Glutamicibacter halophytocola* PFE44 | Coconut fiber | SAMN40757910 |
| *Paenarthrobacter ureafaciens* S54 | Seed | SAMN40757911 |
| *Paenarthrobacter* sp. S56 | Seed | SAMN40757912 |

### 2.2 *In vitro* evaluation of bacterial strains cross-compatibility

The strains were first evaluated *in vitro* by a ‘cross test’ to assess the reciprocal antagonistic activity. The assay was performed on Luria-Bertani Agar (LBA), Tryptic Soy Agar (TSA) and Potato Dextrose Agar (PDA), with three replicates per medium. The plates were incubated at 28°C for 48-96 h. Lack of microbial growth (zone of inhibition) at the intersections was indicative of the antimicrobial activity of the cultures, but the cultures growing in close proximity were compatible to each other. Three bacterial SynComs with different richness levels (4, 6 and 10 strains) were assembled. Each SynCom was tested for strains’ reciprocal compatibility on the same media and growth conditions as above.

### 2.3 Bacterial growth conditions and inoculum preparation

Strains were routinely maintained on Nutrient Agar supplemented with 1% (w/v) dextrose (NDA) at 27 ± 1°C and long-term stored in LB supplemented with 20% (vol/vol) glycerol at -80°C. For the inoculum preparation, all bacterial strains were individually grown in LB broth for 24 h at 27 ± 1°C in a rotary shaker (180 rpm). Bacterial cultures were centrifuged at 5,000 rpm for 15 min, and after discarding the supernatant, the pellets containing the bacterial cells were resuspended in sterile distilled water and the density was normalized to an OD600 of 0.1, containing approximately 10^8^colony forming units (cfu)·mL^-1^. SynComs were assembled mixing bacterial suspensions in equal proportions (1:1:1…).

### 2.4 Plant growth promotion assays

One-month-old seedlings of *Solanum lycopersicum* cv. ‘Proxy’ and cv. ‘Pizzutello’ were produced under standard conditions in a commercial nursery in Ragusa, Italy. The seedlings were individually transplanted in pots (8 cm Ø) filled with a commercial potting substrate (Professional Mix, Vigorplant). After transplanting, 20 mL of the SynCom suspensions (water for the control) were applied in each pot by soil drenching close to the plant crown. Nine and twelve replicates were used for each treatment for ‘Proxy’ and ‘Pizzutello’, respectively. The trial was carried out in a growth chamber under controlled conditions (25°C, RH 60-70%, 16h-light/8h-dark). Plants were monitored regularly and watered on a daily basis. Each pot was rearranged randomly every two-three days. Four weeks after the treatment, the following growth attributes were recorded for both cultivars: plant height, shoots and roots fresh and dry weights. Plant height of ‘Proxy’ seedlings was also measured weekly and rhizosphere samples were collected for microbiome analysis.

### 2.5 Analysis of rhizosphere microbial communities

#### 2.5.1 Experimental design and DNA extraction

Rhizosphere soil from the ‘Proxy’ plants was collected at four time points: T0, beginning of the trial (a few hours after SynComs treatment) and at one (T1), two (T2) and four weeks (T4) post-treatment. At each sampling time, 9 plant per thesis were removed from the pot and the roots were gently shaken to remove excess soil particles. Each sample was formed by five grams of roots with adhering soil from three randomly selected plants thus forming 3 replicates per thesis. Samples were suspended into 20 ml of sterile saline buffer, vortexed and then centrifuged (13,500 rpm, 20 min at 4°C) (Anzalone *et al*., 2022). Pellets were stored at −80°C for subsequent experiments. DNA extraction was performed with the DNeasy PowerSoil Pro Kit (Qiagen, Hilden, Germany) following the manufacturer’s protocol. DNA concentration and purity were determined with a NanoDrop 1000 spectrophotometer (Thermo Scientific, Wilmington, DE, USA) prior to downstream analyses.

#### 2.5.2 16S rRNA and ITS amplicon sequencing

Library preparation and amplicon sequencing were conducted at IGA Technology Services (Udine, Italy). For bacterial community profiling, the V3–V4 hypervariable region of the bacterial 16S rRNA gene was amplified with primers 16S-341F and 16S-805R (Klindworth *et al*., 2013). Peptide nucleic acid (PNA)-clamping was applied during the first 16S rRNA gene amplification step to block amplification of host chloroplast and mitochondrial 16S rRNA gene sequences. For fungal community profiling, primers ITS1 and ITS2 (White *et al*., 1990) were used to amplify part of the ITS1 region of the fungal rRNA operon. The 16S and ITS libraries were sequenced on an Illumina NovaSeq6000 instrument (Illumina, San Diego, CA, USA) using 250-bp paired-end mode. Reads ends were overlapped to generate high-quality full-length sequences and ensure accurate taxonomic classification.

#### 2.5.3 Bioinformatic analysis

Merging forward and reverse reads, quality filtering and trimming, and Amplicon Sequence Variants (ASVs) generation were performed using DADA2 (v. 1.26.0) (Callahan *et al*., 2016) in R (v. 4.0.2) (R Core Team, 2022). The taxonomic assignment of ASVs was performed using the 16S SILVA 138 (Quast *et al*., 2013) database for 16S reads, whereas the UNITE database was considered for ITS reads (version 9.0, all eukaryotic dynamic) (Abarenkov *et al*., 2024). Plant-related (e.g., chloroplast and mitochondria) and unassigned ASVs were filtered out from the 16S ASV table. α- and β-diversity analysis was performed using the phyloseq package (version 3.17) in R (v. 4.0.2). In order to evaluate the diversity within each sample, α diversity was estimated considering the Chao1 richness and Shannon diversity indices. Statistical significance of α diversity was analyzed using the Kruskal– Wallis test. β diversity was assessed through the Bray–Curtis dissimilarity values and depicted with a Principal Coordinate Analysis (PCoA) to evaluate diversity within each group and among the groups of samples. The PERMANOVA test was conducted to assess statistical significance between each group of samples through vegan (Oksanen *et al*., 2007) in R. Differential abundance analysis was performed with DESeq2 (v. 1.40.2) (Love *et al*., 2019) to detect the enriched and depleted bacterial and fungal genera in the SynCom-treated plants compared to the control plants.

Additionally, the PICRUSt2 tool was used to predict the putative metagenome functions based on the ASVs. The genes of the predicted metagenomes were functionally annotated considering the Kyoto Encyclopedia of Genes and Genomes (KEGG) pathways, using ggpicrust2 (Yang *et al*., 2023). The resulting data set was filtered at level 1 to exclude categories not relevant in plant samples (i.e. organismal systems and human diseases). Differential abundance analysis was performed with DESeq2 to assess the enriched and depleted putative functions in the SynCom-treated plants compared to the control plants.

### 2.6 Identification of the inoculated strains in the microbiome

The 16S rRNA gene sequences of the inoculated strains were extracted from the whole genome sequences from Nicotra *et al*. (2024). To cross-reference whether the 16S rRNA gene sequences can be found in microbiome sequences of the rhizosphere of the SynCom-treated plants, they were compared with the 16S rRNA gene amplicon-based metagenomic data of rhizosphere samples using the Basic Local Alignment Search Tool BLASTN (http://www.ncbi.nlm.nih.gov). Sequences with ≥ 99.5% similarity were assigned to the same ASV (Hu *et al*., 2021). This cut of line (≥ 99.5% sequence similarity) was chosen to not overestimate the percentage of matching sequences, due to shorter length of 16S rRNA amplicon sequences as compared to whole 16S rRNA sequence of the inoculated strains. In case of multiple matches, the ASV with the highest identity percentage was selected from those exceeding the set threshold.

### 2.7 Statistical analysis

Data from the PGP experiments were analysed by analysis of variance (ANOVA) using Minitab 20 statistical software (Minitab, Inc., State College, PA). Means were separated using Tukey’s post-hoc HSD test.

## 3. Results

### 3.1 Assembly of bacterial SynComs

In this study we used ten genome-sequenced PGP bacterial strains (Nicotra *et al*., 2024) to evaluate their performance when used as SynComs. Strains identification, source and genome ID are described in Table 1. Strains reciprocal growth inhibition was evaluated *in vitro* by a ‘cross test,’ revealing inhibition events on PDA but not on LBA or TSA (Supplementary table S1). Notably, *Pseudomonas* strains inhibited the growth of all other bacterial strains, regardless of taxon, and partially inhibited each other as well (Supplementary table S1). The only exception was recorded for *Leclercia* sp. S52 that was able to grow in close proximity to *P. simiae* POE78A. *B. velezensis* PFE11 showed inhibition activity against the other *B. velezensis* strains (PFE42 and PSE31B). Zones of inhibition were also detected cross-streaking *Leclercia* sp. S52 against *Paenarthrobacter* strains (S54 and S56), *Chryseobacterium* sp. POE47 and partially against *B. velezensis* PSE31B (Supplementary table S1). Based on this preliminary result, three bacterial consortia were assembled: starting with a four-member SynCom including only *in vitro* compatible bacterial strains, then increasing the richness to 6 by adding the two *Pseudomonas* strains, and finally combining all 10 strains (Table 2).

**Table 2.** Composition of bacterial SynComs assembled in this study.

| Species | Strain | MIX1 | MIX2 | MIX3 |
| --- | --- | --- | --- | --- |
| *Bacillus velezensis* | PFE42 | x | x | x |
| *B. velezensis* | PSE31B | x | x | x |
| *Glutamicibacter halophytocola* | PFE44 | x | x | x |
| *Leclercia* sp. | S52 | x | x | x |
| *Pseudomonas salmasensis* | POE54 |  | x | x |
| *P. simiae* | POE78A |  | x | x |
| *B. velezensis* | PFE11 |  |  | x |
| *Paenarthrobacter ureafaciens* | S54 |  |  | x |
| *Paenarthrobacter* sp. | S56 |  |  | x |
| *Chryseobacterium* sp. | POE47 |  |  | x |

The growth of the bacteria composing each SynCom was evaluated on three different media (Figure 1). On LBA and TSA, the bacterial strains did not inhibit each other, exhibiting overlapping growth between cultures (Figure 1). However, when streaked on PDA, MIX2 and MIX3, both including the *Pseudomonas* strains showed inhibition zones between cultures (Figure 1). Specifically, in MIX2 the inhibition of *G. halophytocola* PFE44 and a reduced growth of the two *Bacillus* strains PSE31B and PFE42 were observed. In MIX3, *G. halophytocola* PFE44 was inhibited, as well as *P. ureafaciens* S54 and *Chryseobacterium* sp. POE47. Similarly to MIX2, the *B. velezensis* strains PSE31B and PFE42 exhibited reduced growth compared to the other tested substrates (Figure 1).

**Fig. 1.**
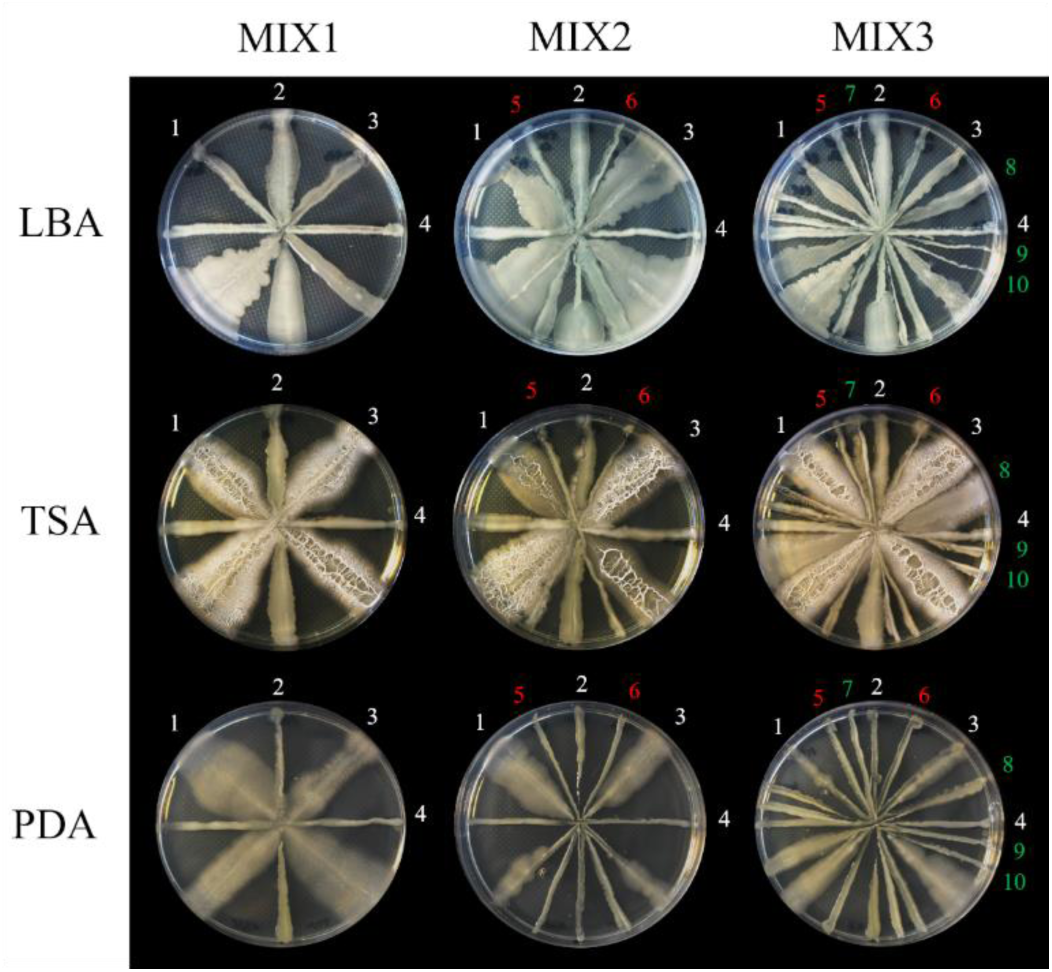
Compatibility assay of the strains in the three SynComs. 1, *B. velezensis* PFE42; 2, *G. halophytocola* PFE44; 3, *B. velezensis* PSE31B; 4, *Leclercia* sp. S52; 5, *P. salmasensis* POE54; 6, *P. simiae* POE78A; 7, *Paenarthrobacter* sp. S56; 8, *B. velezensis* PFE11; 9, *P. ureafaciens* S54; 10, *Chryseobacterium* sp. POE47. Red and green colors indicate the strains progressively added to the MIX1 to obtain MIX2 and MIX3, respectively. LBA, Luria Broth Agar; TSA, Tryptic Soy Agar; PDA, Potato Dextrose Agar.

### 3.2 Effect of SynComs inoculation on tomato growth

Tomato seedlings ‘Pizzutello’ and ‘Proxy’ were treated by soil drenching (T0) with the SynComs consisting of 4 (MIX1), 6 (MIX2), or 10 (MIX3) strains (Table 2). The treatments resulted in a significant increase in plant height four weeks after the treatment (Figure 2). The increase was more pronounced in the indeterminate-growth variety, ‘Proxy’, than in the determinate bush-type ‘Pizzutello’. The growth-promoting effects of the SynComs were further supported by the fresh and dry weight measurements of the shoots, which were significantly higher four weeks after treatments (p < 0.0001 and p = 0.001, respectively) (Figure 3A-B). However, the effect on root biomass was statistically significant only in ‘Pizzutello’ seedlings treated with MIX1 and MIX2, with a partial effect observed in ‘Proxy’ seedlings treated with MIX2 (Figure 3A-B). When considering whole-plant data, all the treatments increased the fresh and dry weight of Pizzutello plants (p < 0.0001), while in Proxy plants MIX2 and MIX3 significantly differed from control ones (p < 0.0001 and p = 0.001, for fresh and dry weight respectively) (Figure 3A-B). Additionally, the height of ‘Proxy’ seedlings was monitored weekly during the sampling for the study of the rhizosphere microbial communities. After one week (T1), all SynCom-treated seedlings showed a significant increase in height compared to control plants drenched with tap water (Supplementary figure S1). The differences between SynCom-treated and control seedlings became even more pronounced from the second week onwards. Seedlings treated with MIX2 and MIX3 exhibited a significant increase in height at all time points compared to both control plants and those treated with MIX1 (p < 0.0001) (Supplementary figure S1). Over the four-week period (T4), plants treated with MIX1, MIX2, and MIX3 demonstrated a 39%, 94%, and 84% increase in height, respectively, compared to the water-treated control seedlings (Supplementary figure S1).

**Fig. 2.**
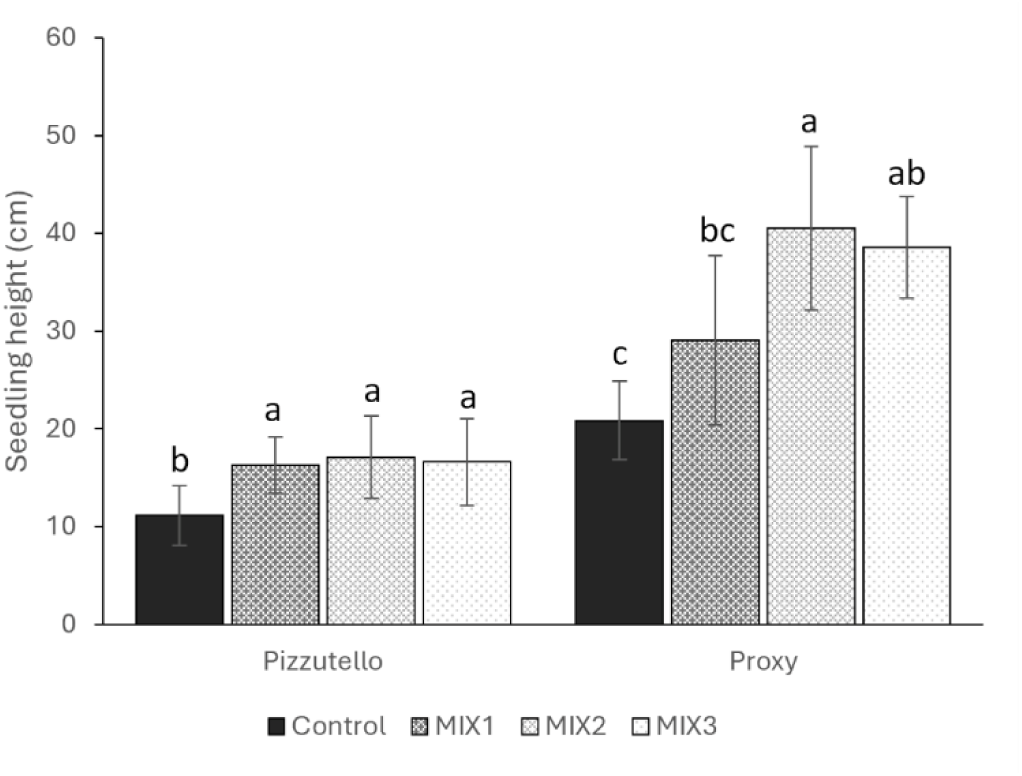
Height of ‘Pizzutello’ and ‘Proxy’ tomato plants four weeks after the soil drenching treatment with the SynComs (water for the control). Different letters denote statistical significance of the values based on post-hoc Tukey HSD test at P = 0.05.

**Fig. 3.**
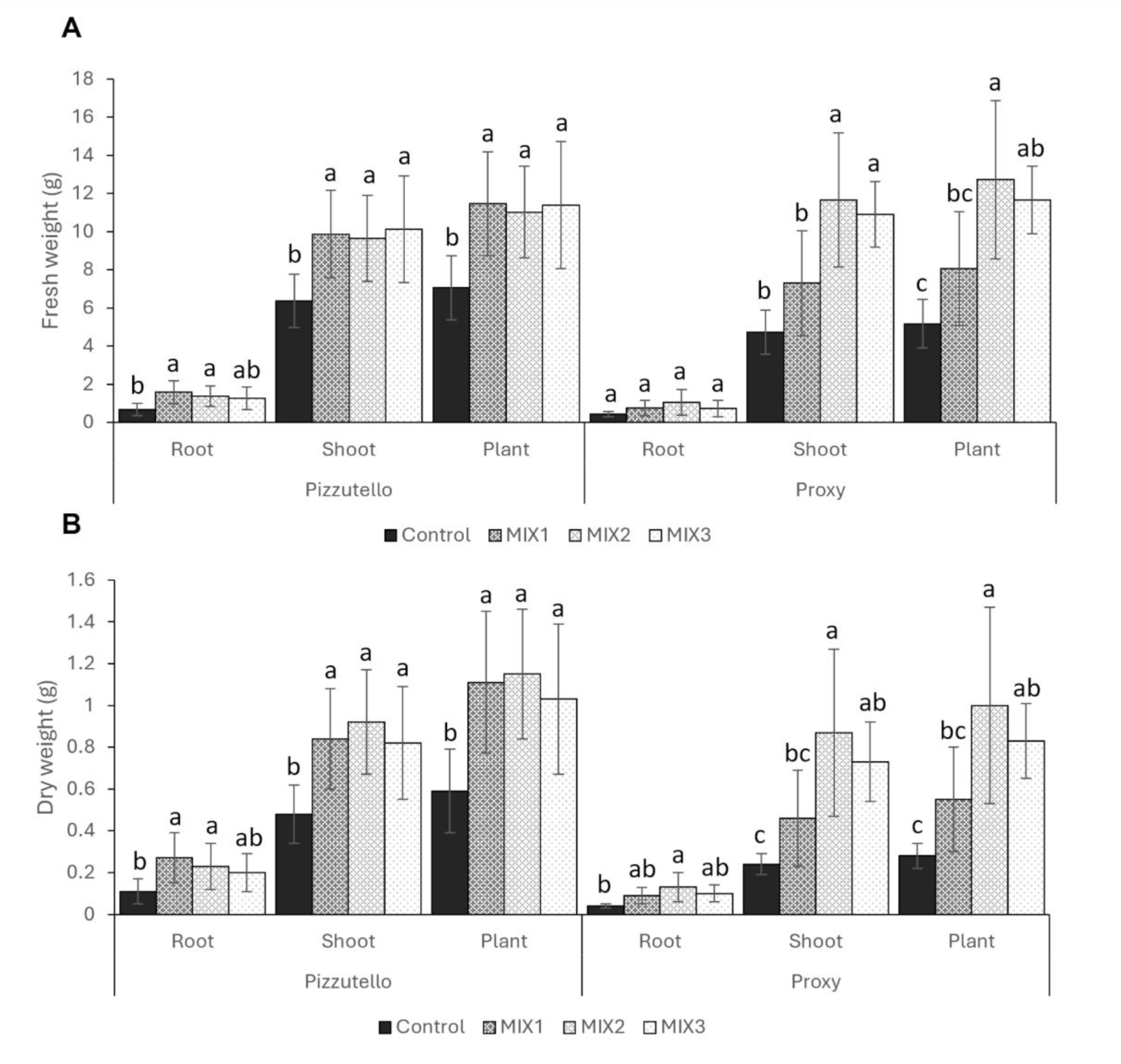
Fresh (A) and dry (B) weight of ‘Pizzutello’ and ‘Proxy’ tomato plants four weeks after the soil drenching treatment with the SynComs (water for the control). Different letters denote statistical significance of the values based on post-hoc Tukey HSD test at P = 0.05.

### 3.3 Effect on resident rhizosphere bacterial and fungal communities

The impact of SynCom treatments on the rhizosphere microbial communities of tomato ’Proxy’ plants was assessed compared to water-treated control plants at four distinct time points: after the bacterial suspension was applied to the soil (T0), and subsequently at one week (T1), two weeks (T2), and four weeks (T4) post-treatment.

Illumina sequencing of the bacterial 16S rRNA gene and the fungal ITS region produced 39,067,796 and 35,164,584 reads, respectively. After paired-end alignments, quality filtering, and deletion of chimeras and singletons, a total of 14,242,196 bacterial 16S reads and 7,623,912 fungal ITS reads were generated from 47 and 48 samples, respectively (one sample was removed from bacterial analysis due to the low number of reads), and assigned to 32,201 bacterial ASVs and 15,182 fungal ASVs.

Two α-diversity indices, Chao1 and Shannon, were calculated for both bacterial and fungal communities across all treatments and time points. Overall, no significant differences in α-diversity were observed between microbial treatments at any of the sampling points, for both bacterial and fungal rhizosphere communities (Figure 4A-B; Supplementary figure S2A-B). However, bacterial communities in control samples exhibited a higher Chao1 index compared to treated samples at all time points, except at the final time point (T4), where the control samples showed the lowest Chao1 value. At this time point, MIX1 and MIX2 treatments displayed the lowest Shannon index (Figure 4A-B).

**Fig. 4.**
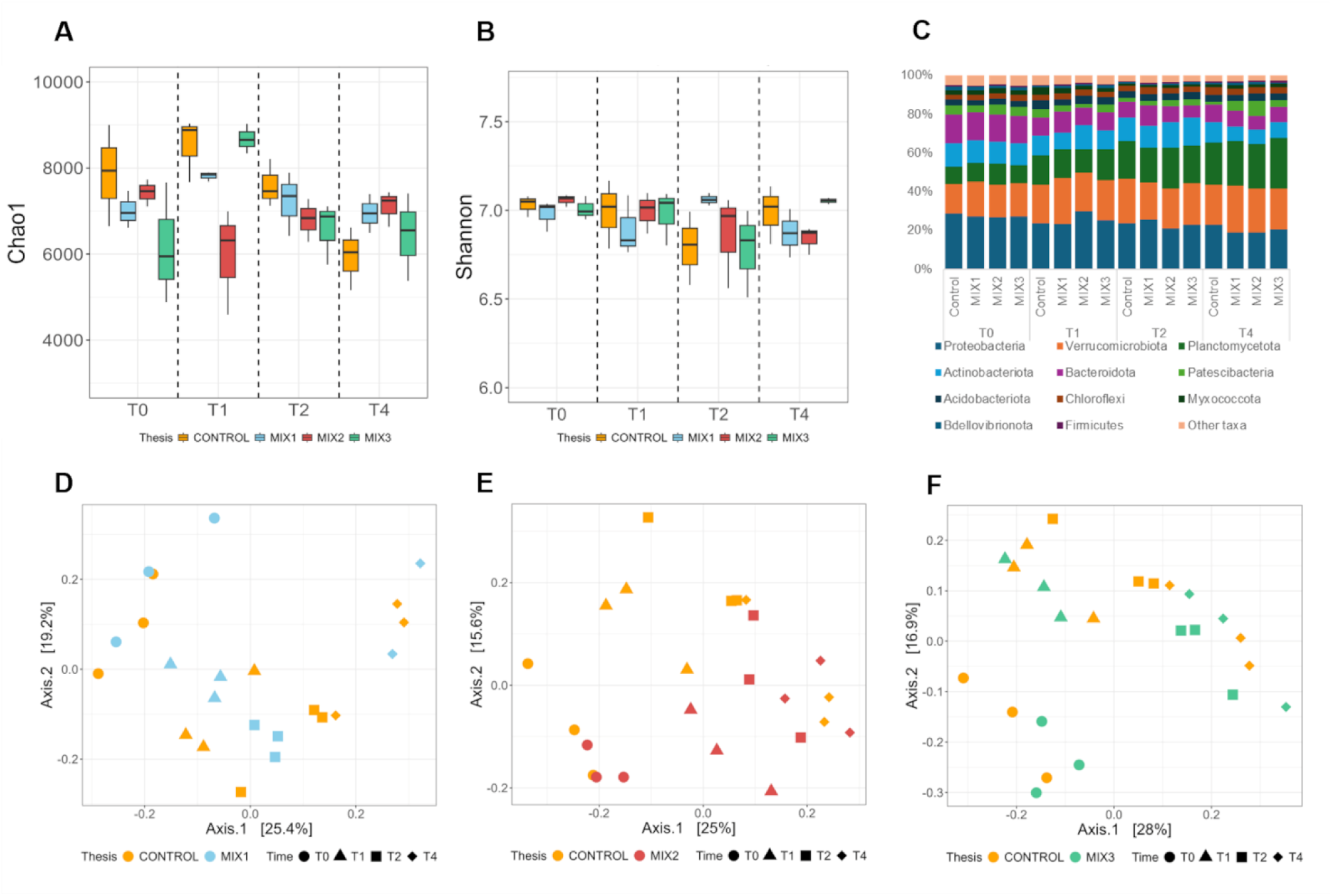
Estimation of the alpha diversity of the microbiome bacterial (A-B) communities in the rhizosphere of control and SynCom-treated tomato plants based on amplicon sequencing data. The observed Chao1 and Shannon indices were used in the alpha diversity analysis. (C) Relative abundances of the bacterial communities at the phylum taxonomic level in the rhizosphere of control and SynCom plants. Taxa less abundant than 1% are reported as “Other taxa”. (D-F) PCoA of rhizosphere bacterial communities in control and SynCom-treated tomato plants based on amplicon sequencing data. Sample clustering was based on the Bray–Curtis dissimilarity matrix. Each point on the graph corresponds to a single sample (biological replication).

For β-diversity, Principal Coordinate Analysis (PCoA) based on Bray–Curtis dissimilarities indicated that time was a major driver of bacterial community diversity, regardless of treatment (Figure 4D-F). However, bacterial communities in MIX2 samples collected one week (T1) and two weeks (T2) after treatment, as well as MIX3 samples at T2, differed from the control (Figure 4E-F). In contrast, neither time nor treatment appeared to significantly influence fungal community composition (Supplementary figure S2D-F).

The relative abundance of bacterial communities at the phylum level reflected the influence of time, as shifts in abundance were observed over the course of the study, despite the overall phylum composition remaining similar. From T0 to T4, a general increase in Verrucomicrobiota and Planctomycetota and a decrease in Proteobacteria, Actinobacteriota, and Bacteroidota was observed across all treatments (Figure 4C).

At T1 the relative abundance of Proteobacteria and Actinobacteriota was higher in the rhizosphere of plant treated with MIX2 than in control plants (29.7% vs 23.4% and 12.4% vs 10.4%, respectively). In addition, the relative abundance of Planctomycetota and Myxococcota was lower in MIX2 compared to the control (12% vs 15% and 1.9% vs 3.7%, respectively) (Figure 4C). At T2 and T4, Patescibacteria abundance was higher in the SynCom treatments compared to the control (from 2.4% to 3.2% vs 1.6% at T2, and from 3.6% to 7.8% vs 1.6% at T4), while Verrucomicrobiota were lower at T2 but higher at T4 in the rhizosphere of SynCom-treated plants (from 19% to 21.7% vs 23% at T2, and from 21% to 24.3% vs 20.4% at T4) (Figure 4C). Additionally, at T4, the relative abundance of Planctomycetota was higher in MIX3 compared to the control (26% vs 21.9%).

Concerning fungal communities, at phylum level tomato rhizosphere was dominated by Ascomycota, followed by Basidiomycota, unidentified Fungi and Mucoromycota (Supplementary figure S2C). The relative abundance of Basidiomicota and unid. Fungi increased while the abundance of Ascomycota and Mucoromycota decreased over time (Supplementary figure S2C).

MIX2 and MIX3 at T2 showed a lower abundance of Ascomycota compared to the control (77.7% and 80.6% vs 84.4%), while in turn Basidiomycota (14% and 10.6% vs 6.3%) and Mucoromycota (4.0% and 5.1% vs 1.8%) were more represented (Supplementary figure S2C). At T4 all the plants treated with the consortia had a higher abundance of Basidiomycota than control (from 10.3% to 11.6% vs 6.4%), and a lower abundance of unid. Fungi (from 6.3% to 11.5% vs 13.7%). In addition, in MIX2 a higher relative abundance of Mucoromycota (2.6% vs from 1.7% to 1.9%) and Fungi_phy_Incertae_sedis (2.6% vs from 0.8% to 1.2%) was observed in the rhizosphere (Supplementary figure S2C).

*Pseudogymnoascus*, belonging to the Ascomycota phylum, was the most represented genus in all samples, with relative abundances ranging from 46% to 65% (data not shown).

### 3.4 Treatments with Syncoms affected more bacterial than fungal communities

Differential abundance analysis was performed to compare the bacterial and fungal genera of the rhizosphere of SynCom-treated plants with the rhizosphere microbial communities of control plants at each time point.

Rhizosphere of plants treated with MIX2 showed the highest number of significant (*p-*value <0.05) differentially abundant bacterial genera compared to the rhizosphere of control plants, except for T2 in which MIX1 had the highest score (Figure 5A). On the overall, in all treatments the number of depleted genera was higher than enriched ones at T0, T2 and T4, with the exception of MIX1 at T2, while at T1 the enriched genera were notably higher than depleted ones in MIX1 and MIX2 (Figure 5A).

**Fig. 5.**
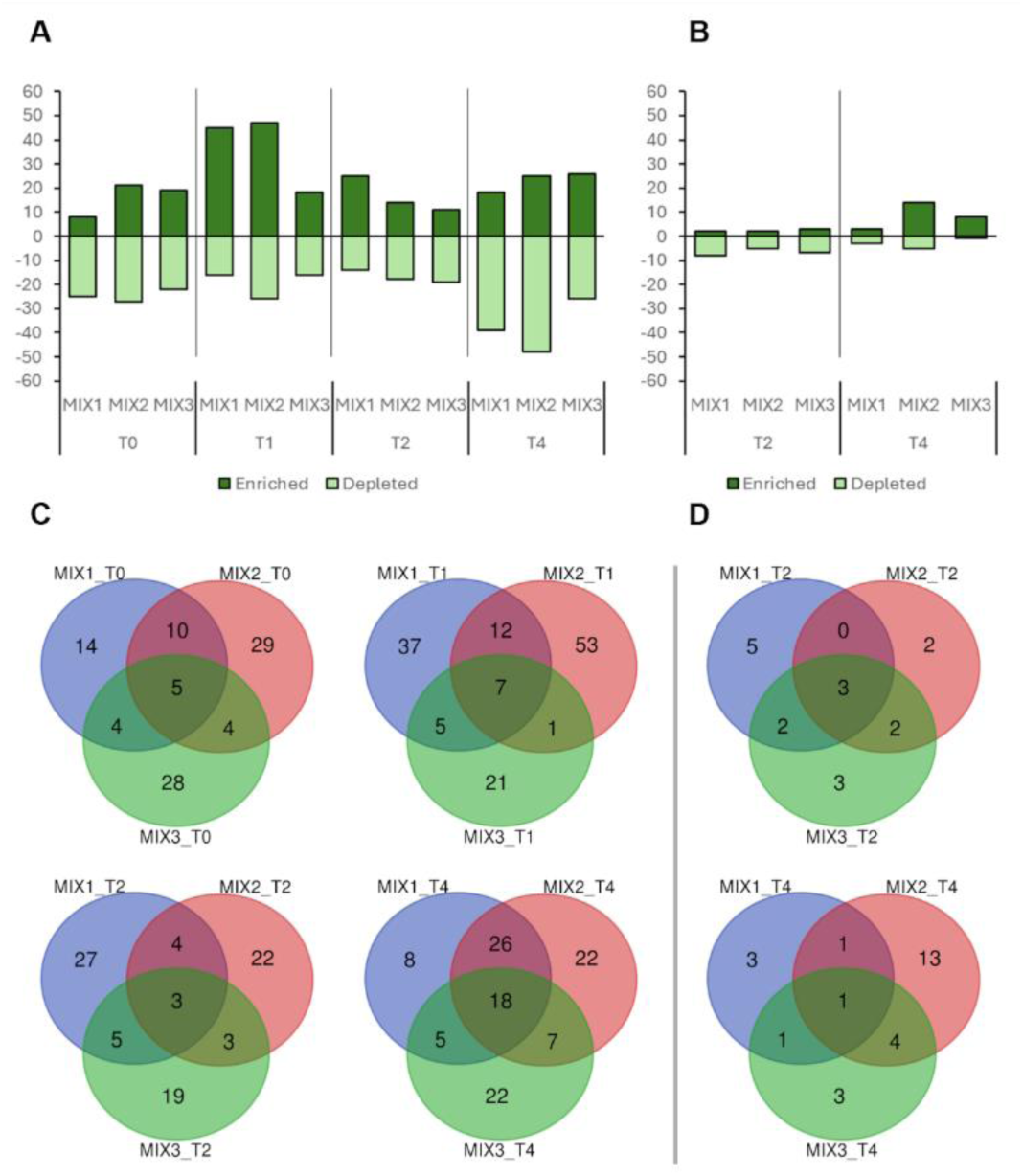
Number of significant (*p-*value <0.05) differentially abundant bacterial (A) and fungal (B) genera present in the rhizosphere samples of plant treated with the SynComs compared to the control at each time point (T0, few hours after SynComs treatment; T1, T2, T4, one, two- and four-weeks post-treatment. Number of unique or shared bacterial (C) and fungal (D) genera in each comparison.

Venn diagrams (Figure 5C) show the number of shared or exclusive bacterial taxa at genus level that had a significant differential abundance (*p-*value <0.05) in the MIX-treated plants compared to the control at each time point. The majority of bacterial genera were unique of a specific MIX at all time points except at T4, in which the number of genera that were in common between all treatments notably increased (Figure 5C). MIX2 and MIX3 showed the highest number of unique genera. In addition, plants treated with MIX1 and MIX2 shared 26 differentially abundant genera, the highest number at the sampling point T4 (Figure 5C).

The differentially abundant bacterial genera that were significantly enriched or depleted (*p-*value <0.05) in at least one comparison at T4 are represented in Figure 6A. Among the 18 differentially abundant genera shared in all treatments, only four were enriched, while the other 14 were depleted compared to control plants (Figure 6A).

**Fig. 6.**
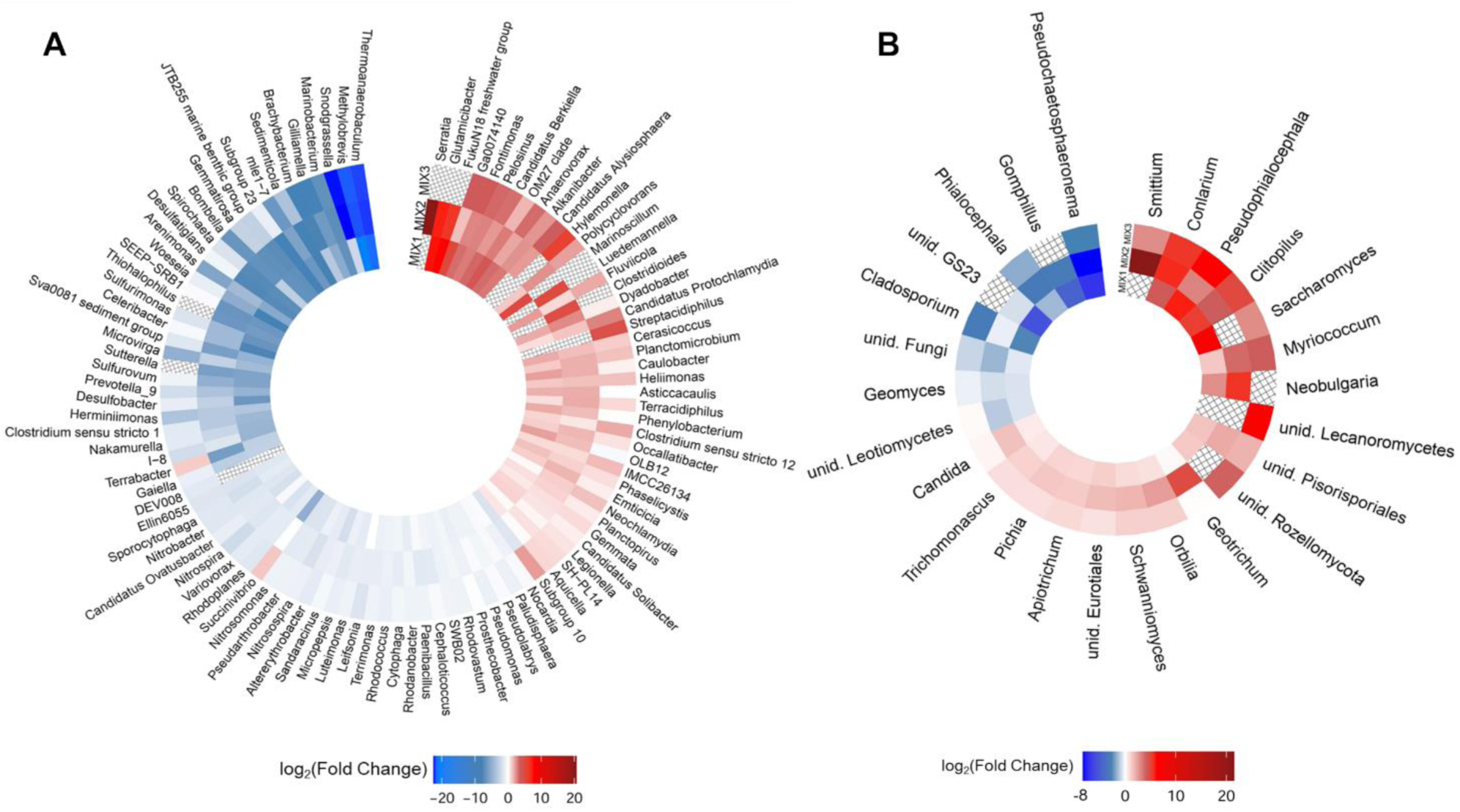
Differentially abundant bacterial genera present in samples treated with SynComs at T4 and significant in at least one comparison (*p-*value <0.05). Each cell depicts the log2 fold change of each bacterial (A) or fungal (B) taxon and is coloured according to depleted (blue), not differentiated (white), and enriched (red) conditions of these samples relative to control samples. Taxa that are not present in a specific sample are denoted with diagonal lines pattern.

Interestingly, most of the differentially abundant genera in the rhizosphere of SynCom-treated plants, compared to the control, are rare taxa, with very low relative abundance in the rhizosphere microbiome (Supplementary figure S3). The only exception was the genus *Gemmata*, which stood out as one of the most abundant bacterial taxa across all treatments and was significantly enriched in the rhizosphere of plants treated with MIX2 and MIX3 (Figure 6A and Supplementary figure S3).

Fungal communities were less affected by SynCom inoculation than bacterial ones. Differentially abundant genera were significantly detected only at two and four weeks after bacterial treatments and were notably lower than the bacterial counterpart (Figure 5B). Interestingly, at T2 the number of depleted genera was higher than enriched genera, while the opposite was observed at T4, with the rhizosphere of plants treated with MIX2 showing a higher number of differentially abundant genera compared to the rhizosphere of control plants (Figure 5D). Only one genus, i.e. *Clitopilus*, was significantly enriched in all treatments compared to control at T4 (Figures 5D and 6B).

### 3.5 Predicted biological functions of rhizosphere bacterial communities

To investigate the potential functional characteristics of the rhizosphere bacterial communities of control and SynCom-treated plants we used the PICRUSt2 tool. The bacterial communities were predicted to participate in various functions, categorized into four major groups encompassing 23 functional subcategories, most of them belonging to metabolic pathways (approximately 71% considering the total genes) (Supplementary figure S4). Genes related to carbohydrate, amino acid and energy metabolism were the most abundant in all treatments, representing approximately 16%, 14% and 9% of the total genes, respectively (Supplementary figure S4). A comparative analysis of the predicted KEGG pathways between the rhizosphere bacterial communities of control and SynCom-treated plants was performed, highlighting significant differences (*p*-value < 0.05, FDR) (Figure 7). The predicted genes of the bacterial communities in the rhizosphere of plants treated with MIX1 were overall enriched compared to control plants, while bacterial communities of plants treated with MIX2 were mostly depleted (Supplementary figure S5). In particular, the depleted gene families in the metabolic pathways were involved in terpenoid and polyketides metabolism as well as metabolism of carbohydrates and amino acids (Figure 7). In turn, an enrichment of genes belonging to the xenobiotics biodegradation and metabolism was found (Figure 7). The opposite trend was observed in the rhizosphere bacterial communities of plants treated with MIX1 (Figure 7).

**Figure 7.**
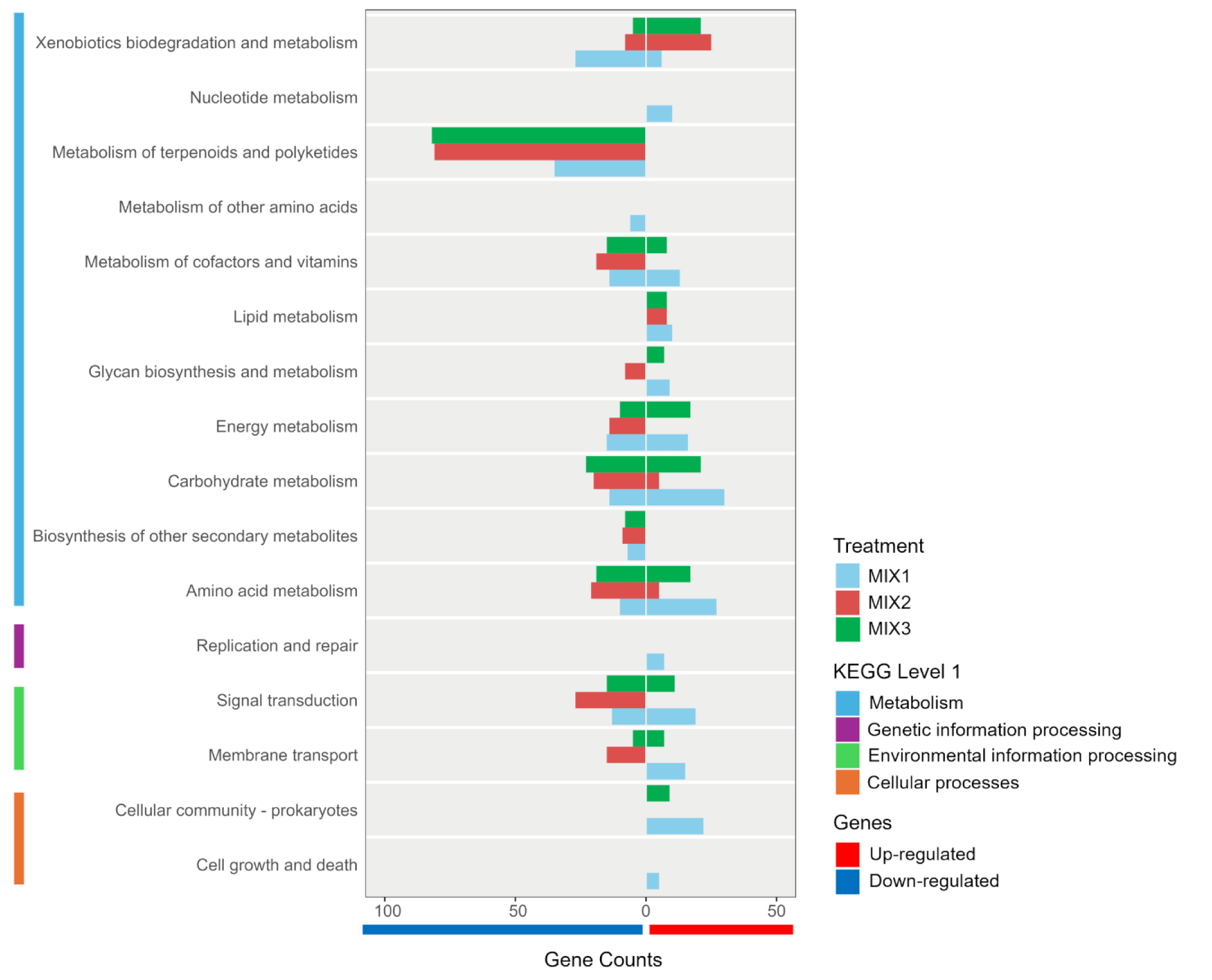
Counts of the predicted functional genes grouped according to KEGG level 1 and 2 categories significantly (*p-*value <0.05, FDR) up- or down-regulated in the rhizosphere bacterial communities of SynCom-treated plants compared to control plants. Only categories with five or more gene counts are represented.

### 3.6 Targeting SynCom strains in the microbiome

The 16S rRNA gene sequences of the inoculated strains in the SynComs were aligned against the 16S rRNA gene amplicon-based metagenomic data of rhizosphere samples. Sequences with ≥ 99.5% similarity were designated targeted ASV sequences according to Hu *et al*. (2021). Eight ASVs with the highest match with the inoculated strains were found and their absolute abundance was employed as a metric to determine the over-time strains abundance in the rhizosphere microbiome (Table 3). The three *Bacillus* strains, all belonging to the same species *B. velezensis* (Nicotra *et al*., 2024), showed the highest sequence similarity with the ASV_407 (Table 3).

**Table 3.** Similarity of the 16S rRNA gene sequences of the 10 bacterial strains with the ASVs of the bacterial communities based on the amplicon sequencing data. The comparison was carried out using the Basic Local Alignment Search Tool BLASTN (http://www.ncbi.nlm.nih.gov). Sequences with ≥ 99.5% similarity were designated targeted ASV sequences according to Hu *et al*. (2021).

| Strain | ASV | Identity (%) |
| --- | --- | --- |
| *Pseudomonas salmasensis* POE54 | 2503 | 100 |
| *Pseudomonas simiae* POE78A | 6620 | 100 |
| *Chryseobacterium* sp. POE47 | 2685 | 100 |
| *Leclercia* sp. S52 | 2430 | 99.73 |
| *Bacillus velezensis* PSE31B | 407 | 99.77 |
| *Bacillus velezensis* PFE42 | 407 | 100 |
| *Bacillus velezensis* PFE11 | 407 | 100 |
| *Glutamicibacter halophytocola* PFE44 | 541 | 99.75 |
| *Paenarthrobacter ureafaciens* S54 | 1004 | 100 |
| *Paenarthrobacter* sp. S56 | 3124 | 100 |

The absolute abundance of the ASVs belonging to the same genera as the inoculated strains decreased over time in the rhizosphere of both control and SynCom-treated plants (Figure 8). On the whole, we were able to detect most of the ASVs of our inoculated strains till the end of the trial, although to a different extent depending on the strain (Figure 8 – Supplementary table S2). In particular, the ASV_2430 matching with the inoculated strain *Leclercia* sp. S52, and included in all SynComs, was not or poorly detected at the end (Figure 8; Supplementary table S2). The same occurred for the ASV_2685 (matching with the strain *Chryseobacterium* sp. POE47) and the ASV_6620 (matching with the strain *P. simiae* POE78A), inoculated in the MIX3 and in both MIX2 and MIX3, respectively (Figure 8; Supplementary table S2).

**Figure 8.**
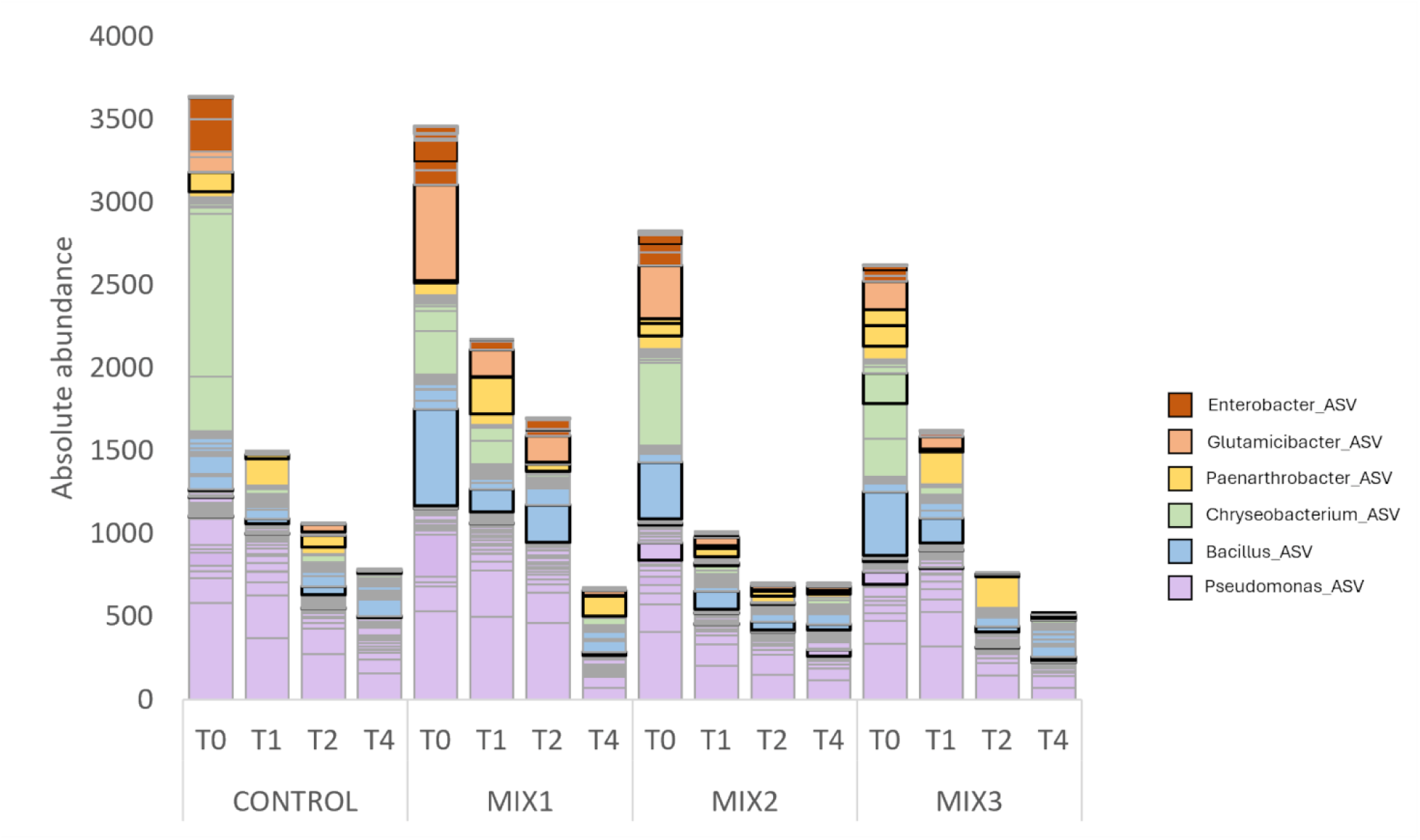
Absolute abundance of the ASVs belonging to the same genera as the inoculated strains in the rhizosphere of tomato plants. Highlighted borders depict ASVs matching with the inoculated strains (≥99.5% sequence similarity). In case of multiple matches, the ASV with the highest identity percentage was selected from those exceeding the set threshold.

Overall, the ASVs of the inoculated strains were detected consistently with the treatments even if in some cases some of the ASVs putatively matching with the inoculated strains (i.e. *Paenarthrobacter* ASV_1004 and *Bacillus* ASV_407) were also detected in the rhizosphere of the untreated control plants, as well as in some treatments where they were not inoculated (Figure 8; Supplementary Table S2).

## 4. Discussion

Three synthetic bacterial communities (SynComs) were assembled from strains isolated from the tomato endosphere, specifically selected as part of the core microbiome of tomato plants in the production chain (Anzalone *et al*., 2022; Nicotra *et al*., 2024). The strains showed plant growth-promoting (PGP) and biological control activities against tomato pathogens (Nicotra *et al*., 2024). Growth chamber assays conducted with two different tomato genotypes revealed that soil drenching with our SynComs significantly enhanced seedling growth. Among the tested SynComs, all containing *Bacillus* strains, MIX2 and MIX3, which also contained *Pseudomonas* species, showed a more pronounced beneficial effect compared to MIX1, which lacked these species. Although bacterial communities were primarily influenced by time, treatment with MIX2 and MIX3 induced shifts in β-diversity compared to control, as early as 1-2 weeks post-treatment. Four weeks after the treatments, changes were still evident with differentially abundant taxa predominantly involving less abundant members and the impacts on the predicted metabolic pathways of the bacterial communities. Despite their low concentrations, most of the SynCom bacteria remained traceable at that time.

Various studies have explored the design of synthetic microbial communities and their impact on plant health and growth. These communities are specifically tailored to interact with plants, promoting growth, enhancing stress tolerance, and boosting disease resistance (Singh *et al*., 2023). SynComs can directly stimulate plant growth through beneficial functions or enhance the activity of specific microbial groups in the rhizosphere by guiding the assembly of resident microbial communities (Hu *et al*., 2021).

However, the selection of SynCom members, their functional properties, and their roles within the community are critical factors for ensuring the effectiveness of SynComs. In this context, core microbiomes represent a promising approach to unraveling the full potential of soil microbiomes to support plant growth (Zhou *et al*., 2024). Native core taxa are expected to colonize soils quickly, persist for extended periods, and contribute to the formation of stable microbial communities (Lemanceau *et al*., 2017; Toju *et al*., 2018). Additionally, the use of multispecies microbial consortia can enhance inoculum survival and establishment, while providing a broader and more diverse range of beneficial functions compared to single-strain inoculants (Hu *et al*., 2021; Martins *et al*., 2023; Singh *et al*., 2023).

The bacterial strains used to assemble the SynComs were selected using a top-down approach matching the data from the microbiome (Anzalone *et al*., 2022) to those of the culturable bacteria obtained from the same samples (Nicotra *et al*., 2024). This set of strains included strains from very well studied biocontrol agents in the genera *Pseudomonas* and *Bacillus* as well as less conventional bacterial genera like *Leclercia*, *Chryseobacterium*, *Glutamicibacter*, and *Paenarthorbacter* (Nicotra *et al*., 2024). Further phenotypic and genomic evidence showed that although shared *in planta* PGPR and biocontrol properties, the *in vitro* antimicrobial activity was actually limited for *Chryseobacterium* sp. POE47 and Micrococcaceae strains (*G. halophytocola* PFE44, *P. ureafaciens* S54 and *Paenarthrobacter* sp. S56 (Nicotra *et al*., 2024). Plant bacterial interaction factor (PIFAR) analysis showed that these strains clustered separately from the others (i.e. *Pseudomonas*, *Bacillus* and *Leclercia*) that showed the highest percentages of toxin-related factors (Nicotra *et al*., 2024).

Treating tomato plants with our SynComs significantly enhanced growth across both tested varieties. In the determinate bush-type variety ’Pizzutello’, all consortia notably increased plant height, fresh and dry biomass. For the indeterminate variety ’Proxy’, only MIX2 and MIX3 improved significantly the growth, with MIX2 having the strongest effect. These enhancements were evident within one week and intensified from the second week onward.

Numerous studies reported on bacterial consortia enhancing tomato plant traits, such as shoot and root length, as well as fresh and dry biomass. In these studies, *Bacillus* strains were combined with other beneficial microbes, such as *Rhizobium* (König *et al*., 2024), *Enterobacter* (Kapadia *et al*., 2021; Ouhaibi Ben Abdeljalil *et al*., 2023), *Achromobacter* (Kapadia *et al*., 2021; Karuppiah *et al*., 2022), *Acinetobacter* (Foughalia *et al*., 2022), and cyanobacteria (Yanti *et al*., 2021), successfully improving tomato growth. Biocontrol activity against tomato plant pathogens (Foughalia *et al*., 2022; Karuppiah *et al*., 2022; Lee *et al*., 2021; Ouhaibi Ben Abdeljalil *et al*., 2023) and abiotic stresses mitigation (Schmitz *et al*., 2022; Wang *et al*., 2016) were also reported.

In our SynComs MIX2 and MIX3, strains of two *Pseudomonas* species, *P. simiae* and *P. salmasensis* were included. Although the *in vitro* antimicrobial activity exerted by *Pseudomonas* against some members of the SynComs, i.e. *G. halophytocola*, *P. ureafaciens*, *Chryseobacterium* sp. and *B. velezensis, in planta* application revealed an improved growth promotion effect compared to MIX1.

Interactions between *Bacillus* and *Pseudomonas* species as PGPR have shown a complex dynamic, often marked by competition in simplified laboratory conditions but potentially stabilizing in diverse natural environments (Lyng *et al*., 2024). However, when in consortia, these species can exhibit synergistic effects, resulting in enhanced biocontrol of pathogens and improved plant growth, particularly through the additive effects of their bioactive compounds and their ability to better colonize plant roots (Lyng and Kovács, 2023). The biocontrol activity against different pathogens like *Cephalosporium* in maize (Ghazy and El-Nahrawy, 2021), *Fusarium* in beans (Kalantari *et al*., 2018), and *Xanthomonas axonopodis* pv. *malvacearum* in cucumber, radish, and cotton (Khabbaz *et al*., 2015; Salaheddin *et al*., 2010) has been associated to the synergy between *Bacillus* and *Pseudomonas* inoculants. In tomato, consortia containing *Bacillus* and *Pseudomonas* species induced systemic resistance against *Sclerotium rolfsii* (Shukla *et al*., 2022). Combination of *Bacillus* and *Pseudomonas* with *Trichoderma* effectively controlled *Fusarium oxysporum* and induced systemic resistance against *Botrytis cinerea* (Minchev *et al*., 2021). Moreover, the synergistic activity of *Bacillus* and *Pseudomonas* isolated from the tomato rhizosphere led to a more robust *in vitro* inhibition of *Alternaria solani* than the single strains (Jia *et al*., 2023).

*Bacillus velezensis* strains (PFE11, PFE42 and PSE31B) and *Pseudomonas simiae* POE78A and *P. salmasensis* POE54 used in this study previously exhibited biocontrol and PGP activities *in vivo* in tomato (Nicotra *et al*., 2024). In particular, *Bacillus* representatives showed the highest growth promotion effect compared to the others with the strain PSE31B being the most effective in controlling Fusarium crown and root rot. *P. simiae* POE78A was the most effective at inducing systemic resistance against bacterial spot caused by *Xanthomonas euvesicatoria* pv. *perforans* (Nicotra *et al*., 2024). Genome analysis highlighted the presence of various beneficial traits related to nutrient acquisition and stress relief. Metabolites predicted by AntiSMASH included surfactin and fengycin lipopeptides in *B. velezensis* among others and obafluorin in *Pseudomonas salmasensis* POE54. PIFAR analysis clustered *Pseudomonas* and *Bacillus* strains together, showing the highest percentages of toxin-related factors compared to the other strains (Nicotra *et al*., 2024).

Studies revealed that *Bacillus*-produced surfactin can promote the colonization of roots by *Pseudomonas* or other beneficial bacteria, allowing them to occupy ecological niches and prevent pathogen invasion (Jia *et al*., 2023; Molina-Santiago *et al*., 2019).

Regarding the other bacterial genera used for designing the SynComs, i.e. *Chryseobacterium* sp., *Paenarthrobacter* sp. and *P. ureafaciens*, there are few reports on members of these genera. *Chryseobacterium balustinum* was used in combination with different species of *Pseudomonas* and cyanobacteria increasing the growth of wheat in a hydroponic growth system (Kholssi *et al*., 2021). Moreover, mixtures of different *Chryseobacterium* species, or *Chryseobacterium* and other PGPR (i.e. *Pseudomonas* sp., *Streptomyces* sp., *Sphingomonas* sp.) protected wheat from *Rhizoctonia solani* and increased the root growth of *Arabidopsis* (Yin *et al*., 2022).

Micrococcaceae family and specifically the genus *Arthrobacter* was reclassified into novel genera, also comprising *Paenarthrobacter* and *Glutamicibacter* (Busse, 2016). Consortia joining *Arthrobacter* sp. with *PaeniBacillus* sp. (Samain *et al*., 2022) or with *Pseudomonas putida* and *Acinetobacter* sp. (Perez *et al*., 2016) were more effective than single-strain inocula in promoting wheat growth and resistance to *Septoria tritici* blotch (STB) and drought stress (Samain *et al*., 2022) and in degrading pesticides in the soil (Perez *et al*., 2016). *Paenarthrobacter nitroguajacolicus* was used in combination with *Pseudomonas*, and *Bacillus* species to accelerate soil recovery in fire-affected soils improving the aggregation of soil particles and the amount of available nutrients (N, P and K) along with the germination and development of the model plant *Bituminaria bituminosa* (Niza Costa *et al*., 2023).

To our knowledge there are no report so far on SynComs comprising *Glutamicibacter* as well as *Leclercia* showing PGP effect in tomato or in other crops. However, considering the numerous beneficial traits identified in the genomes of strains belonging to these genera (i.e. exopolysaccharides, hormones, siderophores) (Nicotra *et al*., 2024) and the activities shown *in planta* (Chen *et al*., 2023; Nicotra *et al*., 2024; Shahzad *et al*., 2017), their inclusion in a consortium could benefit the plant and improve its growth.

Treatment by soil drenching with the bacterial SynComs affected microbial communities of the rhizosphere. Bacterial communities one and two weeks after the treatment with MIX2 and MIX3 according to the beta-diversity were the most influenced. However, several bacterial genera in the communities of the rhizosphere of tomato seedling treated with all the three SynComs were differentially abundant at all sampling times. When introduced microorganisms interact with the resident microflora, they not only directly contribute to plant growth but can also induce indirect effects on the microbial community (Negi *et al*., 2024; Vuolo *et al*., 2022). These indirect effects are mediated by changes in the diversity, composition, and function of the existing microbiome (Castro-Sowinski *et al*., 2007; Vuolo *et al*., 2022). Adding non-native microorganisms to an established microbiome has been demonstrated to significantly impact resident microbial communities in many studies and this change has been correlated to plant health and growth (Chang *et al*., 2024; Hu *et al*., 2021; Sharma *et al*., 2022; J. Wang *et al*., 2021; Zhang *et al*., 2022; Zheng *et al*., 2020). In some cases, the extent of these changes was correlated with the diversity and richness of the introduced consortium, with greater diversity typically leading to more pronounced changes (Hu *et al*., 2021; Rivett *et al*., 2018).

Bacterial community differences between treated and control plants at four weeks after the treatment mostly rely on less abundant bacterial taxa although the plants showed an increase in height from approximately 40% (MIX1) to more than the 80% (MIX2-3).

This suggests the ability of the consortia to rapidly alter the microbial community structure and underscores the importance of selecting SynComs that can establish themselves early in the microbial community, potentially yielding long-term benefits for plant health. Hu *et al*. (2021) reported that the beneficial effects of a *Pseudomonas* consortium on tomato growth were more closely linked to changes in resident community diversity and composition—particularly an increase in the abundance of initially rare taxa—rather than the direct introduction of plant-beneficial traits by the consortia.

Bacterial community dynamics were strongly influenced by the time both in the control and treated samples. A meta-analysis by Shade *et al*. (2013a) showed that archaeal and bacterial communities exhibit significant temporal variability, with the most rapid changes occurring between one day and one month. Richness and evenness of rhizosphere bacterial communities in tomatoes were already reported to be primarily influenced by time (Novello *et al.,* 2024). The effect of time was particularly evident when monitoring the abundances of bacterial genera and even the ASVs corresponding to the SynCom members that severely decrease.

Overall, the inoculated strains were detected till four weeks after the treatments, although they showed a considerable decline compared to the time of inoculation. Moreover, some ASVs were not or poorly detected at the end, in particular those ones matching with *Leclercia* sp. (present in all the SynComs), *Chryseobacterium* sp. and *P. simiae* strains.

The comparison of the ASV/OTU sequences with those of the 16S rRNA gene of the bacterial inoculant was already used by other authors although with different aims and results (Cui *et al*., 2021; Hu *et al*., 2021; Schmitz *et al*., 2022). In general, as expected, the matches between a bacterial strain and a community member were not always unambiguous; therefore, a similar sequence could be detected in the control samples as also observed in other studies (Cui *et al*., 2021; Schmitz *et al*., 2022). Moreover, in our study, multiple *Bacillus* strains were found to be highly homologous to the same ASV. However, it is noteworthy that even just a few hours after soil drenching commercial plantlets in a commercial horticultural substrate with billions of bacterial cells, the introduced bacteria were not strongly represented in the samples, as compared to the substrate microbial communities. Higher relative abundances of SynCom bacterial strains were however observed in other studies in the tomato root environment (Schmitz *et al*. 2022).

Innovative approaches, such as metagenomics applying long-read sequencing to the 16S rRNA gene, other housekeeping genes, or whole-genome shotgun sequencing, could provide more accurate insights for precisely monitoring the fate and impact of introduced microbial strains within a complex community contex(Trivedi *et al*., 2021).

## 5. Conclusions

This study builds upon earlier findings on ten bacterial strains with plant growth-promoting and biocontrol activities against tomato pathogens. Here, these strains were assembled into model bacterial consortia to investigate their effects on tomato growth and the rhizosphere microbiome. The results demonstrate that all consortia positively influenced tomato growth; however, two of the three consortia, characterized by the inclusion of *Pseudomonas* strains, exhibited distinctive effects on the rhizosphere microbial communities. These findings suggest an additive effect of *Pseudomonas* strains *in vivo*, which may enhance their functional impact on plant growth promotion and microbial dynamics.

A detailed analysis of the rhizosphere microbiome revealed that the most significant changes occurred in the less abundant taxa, emphasizing the critical role of these groups in mediating plant-microbe interactions. Such shifts underline the complexity of microbial community dynamics and their potential contributions to plant health and resilience. These observations align with similar findings in other studies, reinforcing the importance of understanding the intricate interplay between introduced bacterial consortia and resident microbial communities.

Overall, this work highlights the potential of bacterial consortia as model systems for exploring the mechanisms underlying plant-microbiome interactions. The observed effects on tomato growth and microbial composition underscore the promise of harnessing microbial consortia for sustainable agricultural practices. Future research should aim to unravel the specific mechanisms driving these interactions and optimize consortium composition to maximize their benefits in diverse environmental and crop conditions.

## CRediT authorship contribution statement

Daniele Nicotra: Conceptualization, Data curation, Formal analysis, Investigation, Methodology, Writing – original draft, Writing – review & editing. Alexandros Mosca: Data curation, Formal analysis, Writing – review & editing, Investigation. Giulio Dimaria: Investigation, Data curation, Formal analysis, Writing – review & editing. Matilde Tessitori: Writing – review & editing.

Ramesh Raju Vetukuri: Conceptualization, Funding acquisition, Project administration, Resources, Supervision, Writing – review & editing, Methodology. Vittoria Catara: Conceptualization, Funding acquisition, Methodology, Project administration, Resources, Supervision, Writing – review & editing.

## Acknowledgments

This article is based upon work from COST Action COST Action MiCropBiomes CA22158, supported by COST (European Cooperation in Science and Technology).

## Appendix A. Supplementary data

The following is the Supplementary data to this article.

## Data availability

Data will be made available on request.

