## Supplementary figures and tables for "Temporal Dynamics of the Tomato Rhizosphere Microbiome in Response to Synthetic Communities of Plant Growth-Promoting Rhizobacteria"

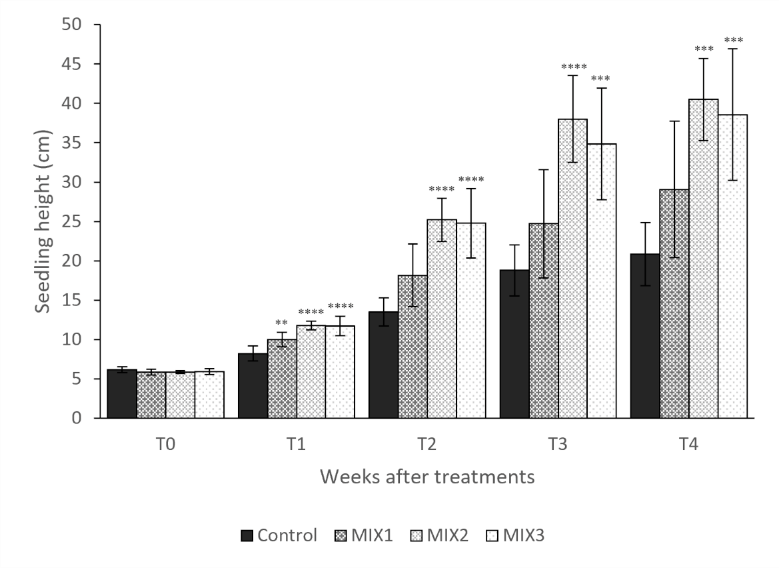

**Figure S1:** Time-course evaluation of tomato seedling height: T0 (few hours after SynCom treatments), T1-4 (1-4 weeks after the treatments). Asterisks denote statistical significance compared to the not treated plants (Control) based on post-hoc Tukey HSD test [**, 0.001≤p≤0.01; ***, 0.0001≤p≤0.001; ****, p<0.0001].

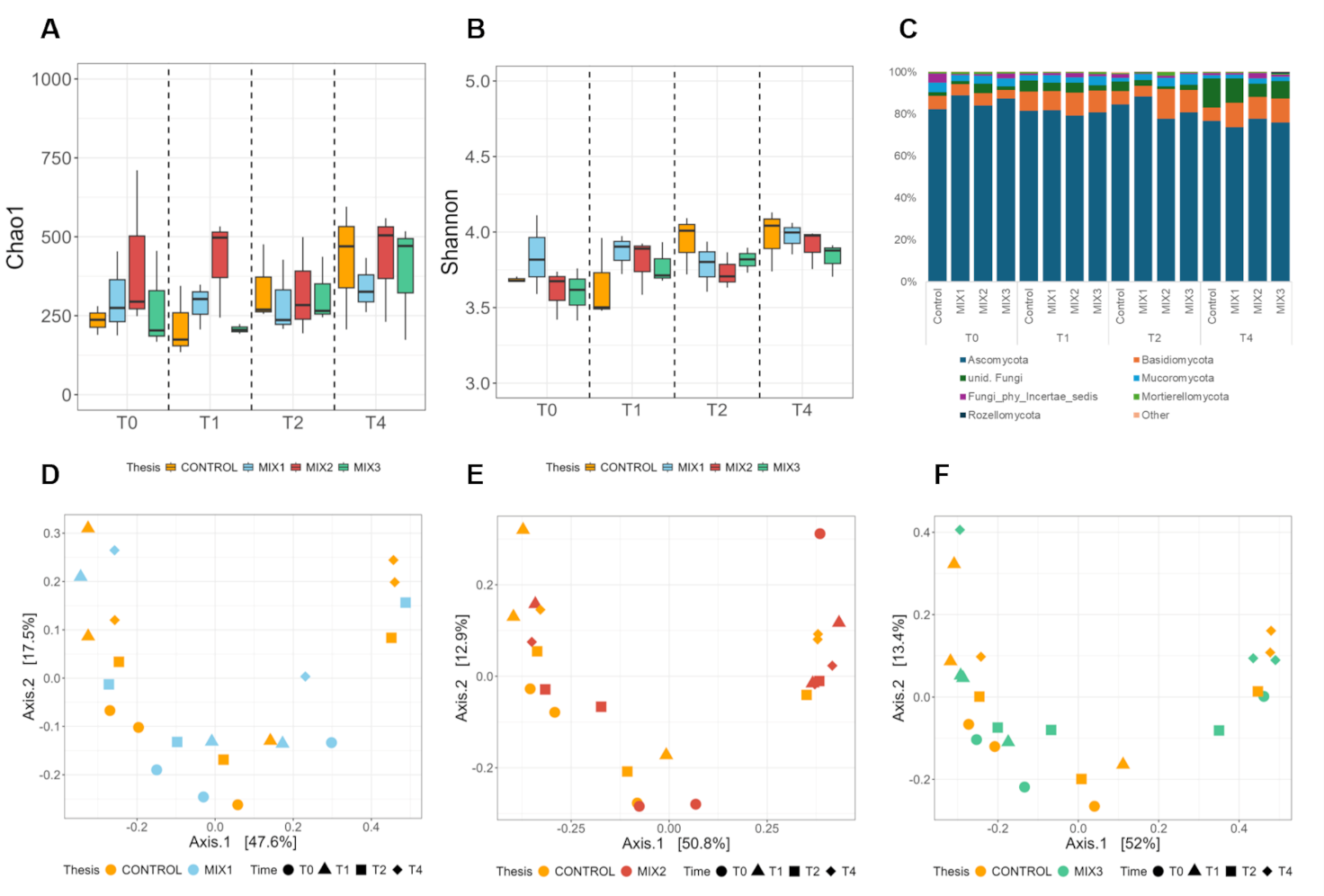

**Figure S2:** Estimation of the alpha diversity of the microbiome fungal (A-B) communities in the rhizosphere of control and SynCom-treated tomato plants based on amplicon sequencing data. The observed Chao1 and Shannon indices were used in the alpha diversity analysis. (C) Relative abundances fungal communities at the phylum taxonomic level in the rhizosphere of control and SynCom plants. Taxa less abundant than 1% are reported as “Other taxa”. (D-F) PCoA of rhizosphere fungal communities in control and SynCom-treated tomato plants based on amplicon sequencing data. Sample clustering was based on the Bray–Curtis dissimilarity matrix. Each point on the graph corresponds to a single sample (biological replication).

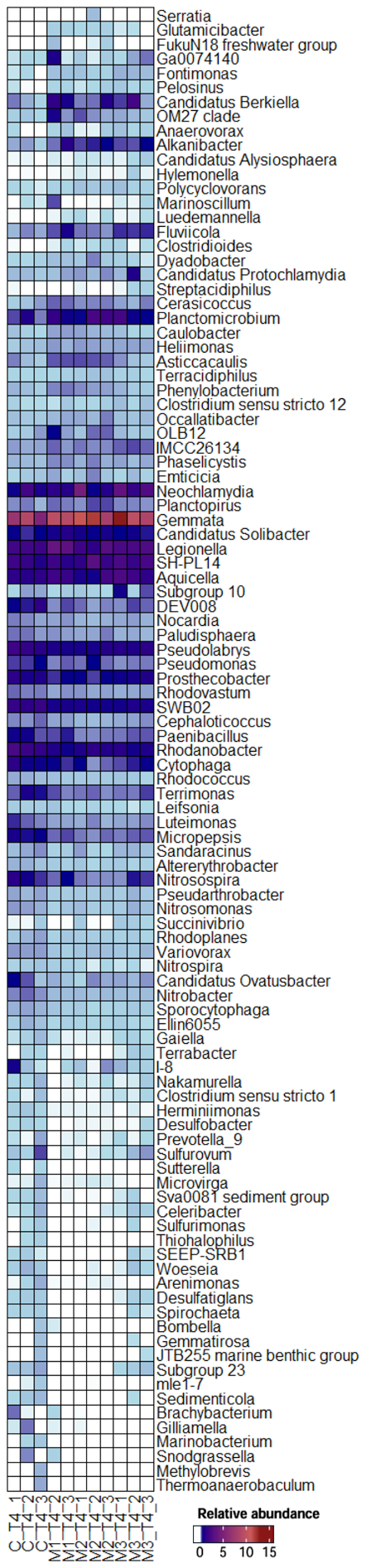

**Figure S3:** Relative abundance of differentially abundant bacterial genera (p-value < 0.05) in the rhizosphere of SynCom-treated plants compared to control at T4. Bacterial genera are indicated in the right; samples are indicated on the bottom. Colors from light blue to dark red represent the lowest and highest relative abundances, respectively. White color represents no abundance of a genus in specific samples.

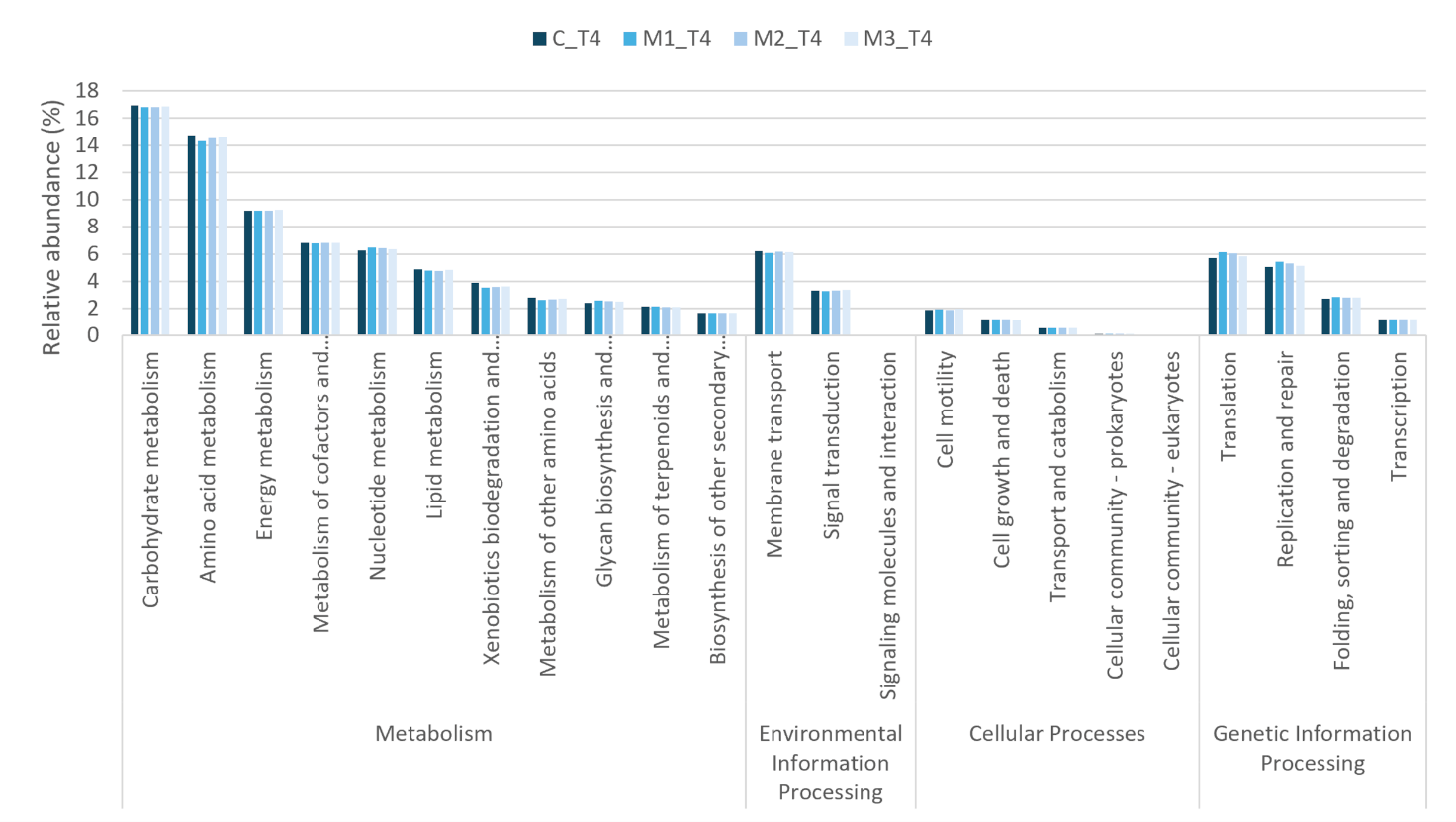

**Figure S4:** Relative abundance of the predicted functions grouped according KEGG level 1 and 2 categories of the rhizosphere bacterial communities of the Control and SynCom-treated plants.

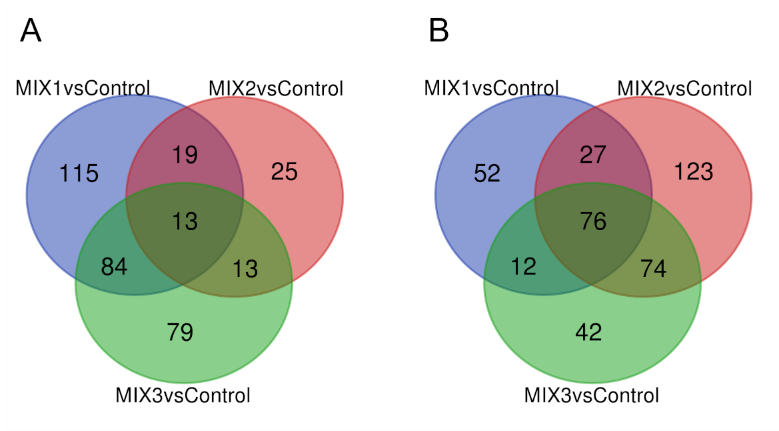

**Figure S5:** Number of exclusive or shared predicted genes significantly (*p*-value < 0.05, FDR) up- (A) or down-regulated (B) in the rhizosphere bacterial communities of SynCom-treated plants compared to control plants.

**Table S1:** *In vitro* reciprocal growth inhibition of bacterial strains evaluated on PDA. 0, no inhibition; 1, inhibition; 2, partial inhibition.

|  | *Bacillus* *velezensis* PFE11 | *Bacillus* *velezensis* PFE42 | *Bacillus* *velezensis* PSE31B | *Pseudomonas* *salmasensis* POE54 | *Pseudomonas* *simiae* POE78A | *Leclercia* sp. S52 | *Chryseobacterium* sp. POE47 | *Glutamicibacter* *halophytocola* PFE44 | *Paenarthrobacter* *ureafaciens* S54 | *Paenarthrobacter* sp. S56 |
| --- | --- | --- | --- | --- | --- | --- | --- | --- | --- | --- |
| *Bacillus* *velezensis* PFE11 | 0 | 1 | 1 | 1 | 1 | 0 | 0 | 0 | 0 | 0 |
| *Bacillus* *velezensis* PFE42 | 1 | 0 | 0 | 1 | 1 | 2 | 0 | 0 | 0 | 0 |
| *Bacillus* *velezensis* PSE31B | 1 | 0 | 0 | 1 | 1 | 0 | 0 | 0 | 0 | 0 |
| *Pseudomonas* *salmasensis* POE54 | 1 | 1 | 1 | 0 | 2 | 1 | 1 | 1 | 1 | 1 |
| *Pseudomonas* *simiae* POE78A | 1 | 1 | 1 | 2 | 0 | 0 | 1 | 1 | 1 | 1 |
| *Leclercia* sp. S52 | 0 | 2 | 0 | 1 | 0 | 0 | 1 | 0 | 1 | 1 |
| *Chryseobacterium* sp. POE47 | 0 | 0 | 0 | 1 | 1 | 1 | 0 | 0 | 0 | 0 |
| *Glutamicibacter* *halophytocola* PFE44 | 0 | 0 | 0 | 1 | 1 | 0 | 0 | 0 | 0 | 0 |
| *Paenarthrobacter* *ureafaciens* S54 | 0 | 0 | 0 | 1 | 1 | 1 | 0 | 0 | 0 | 0 |
| *Paenarthrobacter* sp. S56 | 0 | 0 | 0 | 1 | 1 | 1 | 0 | 0 | 0 | 0 |

**Table S2:** Absolute abundance (number of reads) of the ASVs matching with the inoculated strains in the rhizosphere of the tomato plants treated with the Syncoms. T0, few hours after SynCom treatment; T1-4 one to four weeks after SynCom treatment.

|  |  | **CONTROL** | | | | **MIX1** | | | | **MIX2** | | | | **MIX3** | | | |
| --- | --- | --- | --- | --- | --- | --- | --- | --- | --- | --- | --- | --- | --- | --- | --- | --- | --- |
| **STRAIN** | **ASV** | **T0** | **T1** | **T2** | **T4** | **T0** | **T1** | **T2** | **T4** | **T0** | **T1** | **T2** | **T4** | **T0** | **T1** | **T2** | **T4** |
| *Bacillus velezensis* PFE11, PFE42, PSE31B | 407 | 11.67 | 31.33 | 50.00 | 8.00 | 581.67 | 137.67 | 226.00 | 24.00 | 344.67 | 107.67 | 52.00 | 30.67 | 383.00 | 150.00 | 33.00 | 19.00 |
| *Glutamicibacter halophytocola* PFE44 | 541 | 0.00 | 0.00 | 44.00 | 0.00 | 579.00 | 163.67 | 161.33 | 28.00 | 318.00 | 47.33 | 23.00 | 24.67 | 172.00 | 71.33 | 12.33 | 7.00 |
| *Paenarthrobacter ureafaciens* S54 | 1004 | 80.67 | 25.67 | 71.67 | 10.33 | 1.67 | 220.00 | 42.00 | 120.00 | 72.00 | 51.67 | 27.67 | 16.00 | 125.33 | 3.67 | 0.67 | 15.00 |
| *Pseudomonas salmasensis* POE54 | 2503 | 0.33 | 0.00 | 0.33 | 0.00 | 0.00 | 0.00 | 0.00 | 0.00 | 105.67 | 1.33 | 0.00 | 37.67 | 76.33 | 12.00 | 3.33 | 0.00 |
| *Chryseobacterium* sp. POE47 | 2685 | 0.00 | 0.00 | 0.00 | 0.00 | 0.00 | 0.00 | 0.00 | 0.00 | 0.00 | 18.00 | 5.67 | 0.00 | 185.67 | 0.00 | 0.00 | 3.00 |
| *Paenarthrobacter* sp. S56 | 3124 | 1.67 | 0.00 | 0.00 | 0.00 | 4.67 | 1.00 | 0.00 | 1.50 | 30.67 | 14.67 | 9.00 | 0.00 | 94.00 | 10.33 | 0.00 | 1.33 |
| *Leclercia* sp. S52 | 2430 | 4.00 | 1.00 | 0.33 | 0.00 | 126.67 | 3.00 | 7.67 | 0.00 | 54.33 | 12.33 | 0.67 | 1.00 | 24.67 | 5.33 | 5.33 | 1.00 |
| *Pseudomonas simiae* POE78A | 6620 | 3.00 | 0.33 | 0.00 | 0.00 | 0.67 | 1.00 | 0.00 | 0.00 | 13.33 | 1.00 | 0.67 | 0.00 | 15.67 | 3.33 | 0.00 | 0.67 |
